# Detection of *Wolbachia* in Field-Collected Mosquito Vector, *Aedes aegypti*

**DOI:** 10.1101/408856

**Authors:** Thaddeus M. Carvajal, Kazuki Hashimoto, Reza Kurniawan Harnandika, Divina M Amalin, Kozo Watanabe

## Abstract

It was the impression from past literature that *Wolbachia* is not naturally found in *Ae. aegypti*. However, there are have been reports that recently reveals the presence of this endosymbiont in this mosquito vector. With this, our study presents additional support of *Wolbachia* infection in *Ae. aegypti* by screening field-collected adult mosquitoes using *Wolbachia*specific 16S rDNA and its surface protein (*wsp*) makers under optimized PCR conditions. From a total of 672 *Ae. aegpyti* adult mosquito samples collected in Metropolitan Manila, Philippines, 113 (16.8%) and 89 (13.2%) individual mosquito samples were determined to be *Wolbachia* infected using the *wsp* and 16S rDNA markers, respectively. The *Ae. aegpyti wsp* sample sequences were similar or identical to five known *Wolbachia* strains belonging to supergroups A or B while majority of 16S rDNA sample sequences were similar to strains belonging to supergroup B. Overall, 80 (11.90%) individual mosquito samples revealed to show positive amplifications in both markers and 69.0% showed congruence in supergroup identification (supergroup B). Our findings illustrate that the infection status of *Wolbachia* in *Ae. aegypti* may appear common than previously recognized.

## INTRODUCTION

Mosquitoes are considered to be medically important insects because of their capacity to carry notable human disease pathogens^1^. Among the known mosquito vectors, *Aedes aegypti* is an efficient and dangerous mosquito vector because of its ability to carry significant arboviral diseases such as Dengue, Chikungunya, Yellow Fever and Zika^2,3^. Despite the development of vaccines, these arboviral diseases are considered to be the leading cause of global disease burden^4^and thus, targeting the mosquito vector is deemed to be the primary control and prevention. A considerable number of vector control strategies had been implemented, but the disease burden continues to increase. Novel and newer approaches are being developed that shows promising outcomes in vector and disease control and one of which is the utilization of the intracellular bacterial endosymbiont, *Wolbachia*^*7-9*^.

*Wolbachia* is a naturally occurring endosymbiont which can be maternally inherited and cause different reproductive alterations in its host to increase their transmission to the next generation^10-12^. In insects, it is estimated to be naturally present in 60-65% of known species^13^. As to date, there are 17 identified major clades or supergroups (A-Q) where a majority are known to infect arthropods such as insects, arachnids, and crustaceans^14^. The pathogenic effects of *Wolbachia* in its host are well-studied and determined to cause sperm-egg incompatibility, parthenogenesis, cytoplasmic incompatibility, and feminization^11,15^. Therefore, utilizing these effects towards medically-important mosquito vectors, such as *Ae. aegypti*, has taken great research strides in the past two decades. The discovery of a virulent *Wolbachia* strain (*w*MelPop) in Drosophila melanogaster was successfully transferred to *Ae. aegypti* where it reduced the lifespan of the mosquito vector^16-18^. In addition to this, *w*MelPop and other *Wolbachia* strains (e.g., *w*Mel) were able to demonstrate conferring resistance on a wide range of insect viruses, especially to human viral pathogens, such as dengue and chikungunya^19-22^. The life-shortening capability plus pathogen interference of this *Wolbachia* strain opened an avenue for its potential use as a biological control agent approach against mosquito-borne diseases. The World Mosquito Program (https://www.worldmosquitoprogram.org), formerly known as the Eliminate Dengue Project, was able to generate stable *Wolbachia*-infected *Ae. aegypti* lines that possessed the ability of pathogen interference from dengue viruses under laboratory conditions. These *Wolbachia* strains showed maternal transmission rates close to 100% and induced high levels of cytoplasmic incompatibility to *Ae. aegypti*^16^. Semi-field cage experiments were also conducted to assess the fitness cost effect of the discovered strain towards the mosquito vector and its ability of these strains to invade the mosquito population. These experiments demonstrated the true potential of the endosymbiont because of the reduced fecundity of *Wolbachia*-infected *Ae. aegypti* as compared to the uninfected wildtype^22^. Australia became the first country to release these *Wolbachia*-infected *Ae. aegypti* into the wild population where it exhibited promising results^23,24^. As to date, this methodological strategy against the mosquito vector, *Ae. aegypti*, is now being tested in eight dengue-endemic countries such as Indonesia, Vietnam, Colombia, and Brazil (https://www.worldmosquitoprogram.org). It claimed this approach is considered to be cost-effective and safer for the environment than conventional insecticide-based measures^19,25^.

With the recognition of about 65% of known insects to be naturally infected with *Wolbachia* including those mosquito species from the genera of *Aedes, Culex, Mansonia*, major mosquito vectors of diseases such as *Ae. aegypti* and Anopheline mosquitoes were reported not to possess this endosymbiont^26-31^. It led to the belief that the presence of *Wolbachia* endosymbiont could be the reason why many of the mosquito species are considered to be weak vectors^23^. Nonetheless, more recent studies show evidence that *Wolbachia* infection in *Ae. aegypti* and *Anopheles gambiae* may appear to be more common than it was previously recognized. Natural *Wolbachia* infections have now been reported in adult, larvae and egg populations of *An. gambiae*^32-34^.

Lately, studies have reported detecting *Wolbachia* from field-collected *Ae. aegypti* samples using either *wsp* marker^35^or 16S metabarcoding^36-37^. Though these studies are commendable, there were still uncertainties in establishing whether the mosquito vector does harbor naturally the endosymbiont. Although metabarcoding studies had a substantial sample size (n=85-270), there were unable to report an accurate estimate of the infection rate because mosquito adult or larval samples were pooled from each location. In contrast, *wsp* detection in *Ae. aegypti* larval samples^35^were screened individually, thus, was able to report the infection rate (50.0%). However, it was difficult to affirm or ascertain its true prevalence since the sample size was small (n=16 individuals). Moreover, there is possibility of a potential bias in reporting a high infection rate if larval samples were collected from the same water container due to the sampling of mosquito siblings from the same female mosquito. Nevertheless, these studies further suggest the likelihood of *Wolbachia* to be naturally associated with *Ae. aegypti*, thus, opening an avenue to re-visit or re-examine its infection status.

Our study aims to present additional support of *Wolbachia* infection found from field-collected *Ae. aegypti* adult mosquitoes using *Wolbachia*-specific 16S rDNA and the *Wolbachia* surface protein (*wsp*) markers. Based on the limitations presented from previous studies, two considerations were applied in addressing these gaps. First, *Wolbachia* screening was done over a large sample size (n=672) and used an individual-based detection of adult *Ae. aegypti* mosquitoes to gain a better estimate of its prevalence in this mosquito vector. Secondly, two molecular markers were used to confirm the detection status and infer the type of *Wolbachia* strains found in *Ae aegypti*.

## METHODS

### Study area and Mosquito collection

The study area is the National Capital Region of the Philippines or also known as Metropolitan Manila. Located on the Eastern shore of Manila Bay in Southwestern Luzon Island (14°50′ N Latitude, 121°E Longitude), it is considered to be one of the highly urbanized and densely populated areas in the Philippines. Dengue disease is endemic in this region where it accounts for 15%-25% of the total number of reported Dengue cases annually in 2009 2014^38^. Vector control programs are being implemented in various localities of the region. Insecticide application and cleaning of the surroundings have been extensively used however its effectiveness is in question because of the constant and unchanging burden of the disease. As to date, the Philippines, especially Metropolitan Manila, has never conducted any *Wolbachia*-based program against *Ae. aegypti*.

Adult mosquito samples were collected using a commercial branded mosquito UVlight trap (Jocanima^®^) installed in the outdoor premises of 138 residential households (sampling sites) from May 2014 – January 2015 (Figure 1a). Collected samples were then sorted and identified as *Ae. aegypti* using available keys^39^. Each sample was then placed in a tube with 99.5% ethanol for preservation. A total of 672 *Ae. aegypti* adult mosquito samples were collected, identified, labeled accordingly (See Supplementary Table 1) and stored at 20°C for subsequent processing.

**Fig 1.**
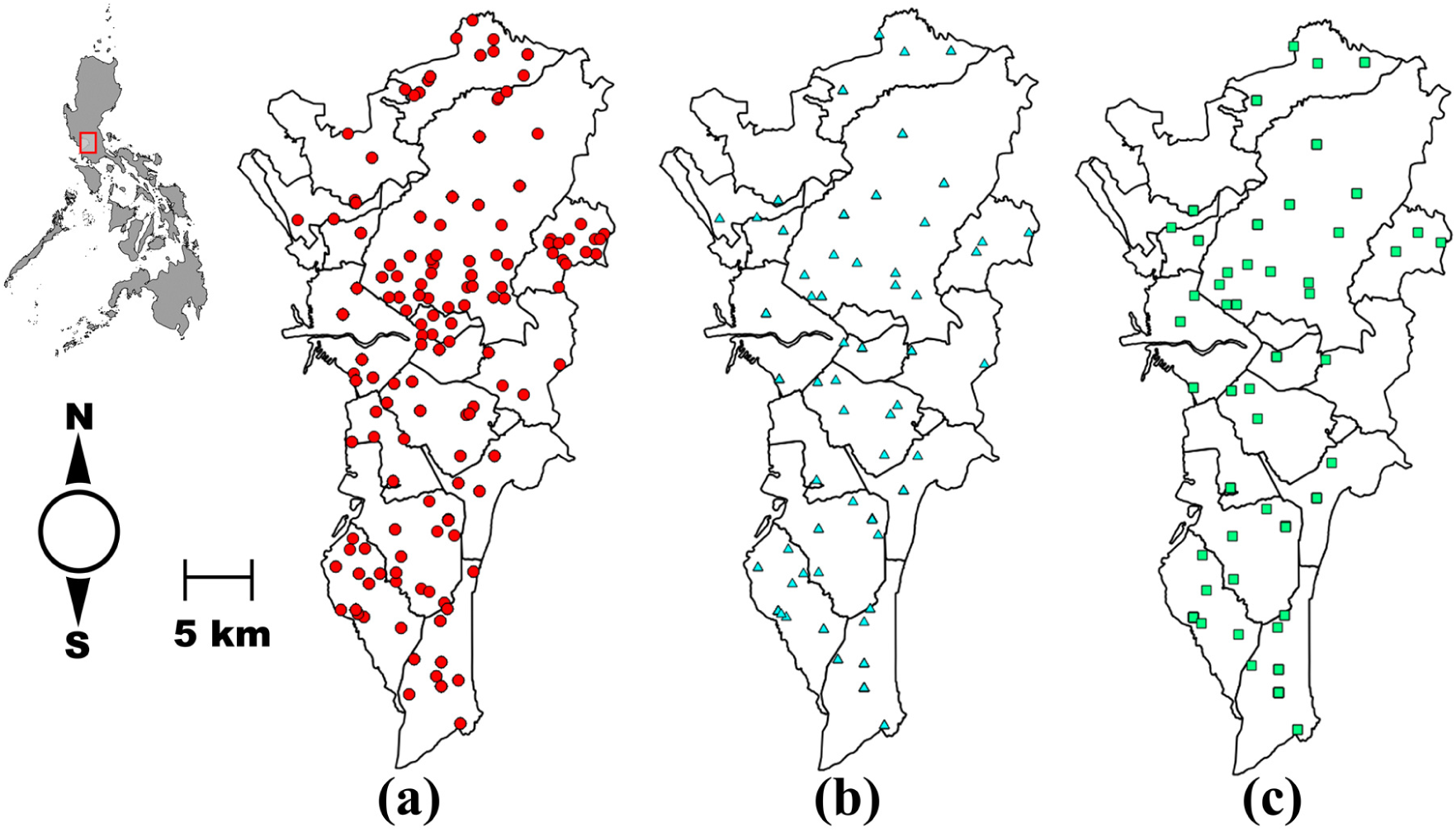
(a) Spatial distribution of the sampling sites (n=138) for collecting adult *Ae. aegypti*. *Wolbachia*-positive sampling sites based on *wsp* (b) and 16S rDNA (c). Details of the number of *Wolbachia*-positive mosquitoes per sampling site is found in the Table S1.

### DNA Extraction, Polymerase Chain Reaction, and Sequencing

Total genomic DNA of each mosquito individual was extracted using the QIAGEN Blood and Tissue DNEasy Kit© following a modified protocol^40^. Our study used two molecular markers for detecting *Wolbachia* infection namely; *wsp*^41^ and 16S rDNA^42^. The primer sequences are as follows: *wsp* 81F (5′TGG TCC AAT AAG TGA TGA AGA AAC) and *wsp* 691R (5′ AAA AAT TAA ACG CTA CTC CA) for *wsp* marker while *Wspecf* (AGC TTC GAG TGA AAC CAA TTC) and *Wspecr* (GAA GAT AAT GAC GGT ACT CAC) for 16S rDNA.

For *wsp* gene amplification, we followed the standard *wsp* protocol^30^where the suggested annealing temperature and number of cycles were 55 °C and 30 cycles respectively. In order to conduct an individual-based detection, we initially performed this protocol in *Culex quinquefasciatus* as our positive control. Certain modifications were made in the standard protocol based on the results where the annealing temperature was set to 57 °C and the number of cycles increased to 35 cycles. This initial modified protocol was performed in individual *Ae. aegypti* samples where it yielded positive faint bands. It prompted us to remodify again the protocol where the annealing temperature is set at 59 °C with 40 cycles and the addition of 10% DMSO (Sigma-Aldrich^®^) that led to desirable results necessary for sequencing. Therefore, a 10 *μl* final reaction volume was used and composed of 10X buffer (TAKARA^®^), 25 mM MgCl_2_, 10 mM of each dNTPs, 10 *μ* M forward and reverse primers, 10% DMSO (Sigma-Aldrich^®^) and 5.0U/ *μl* of *Taq* DNA polymerase (TAKARA^®^). The final thermal profile consisted an initial denaturation of 95 °C for 3 minutes, followed by another denaturation temperature of 95 °C for 1 minute, an annealing temperature of 59 °C for 1 minute and an extension temperature of 72 °C for 1 minute for 40 cycles, and accompanied by a final extension temperature at 72 °C for 3 minutes.

On the other hand, 16S rDNA gene amplification used a 10 *μl* final reaction volume and composed of 10X buffer (TAKARA), 25 mM MgCl_2_, 10 mM of each dNTPs, 10 *μ*M forward and reverse primers, 10% DMSO (Sigma-Aldrich^®^) and 5.0U/ *μ*L of *Taq* DNA polymerase (TAKARA^®^). Thermal profiles follow the protocol of Simões et al^42^with initial denaturation temperature at 95 °C for 2 minutes, followed by two cycles of 95 °C for 2 minutes of denaturation, annealing temperature of 60 °C for 1 minutes and extension temperature of 72 °C for 1 minute, afterwards 35 cycles of denaturation of 95 °C for 30 seconds, annealing temperature of 60 °C for 1 minute and extension temperature of 72 °C for45 seconds and final extension at 72 °C for 10 minutes.

All PCR amplification experiments included positive and negative controls. The positive control is a *Wolbachia*-infected *Cu. quinquefasciatus* sample while the negative control consisted of water as the template. The product size of each molecular marker was checked through electrophoresis with 1.5% agarose gel set at 100 volts for 30 minutes. The size of the amplified *wsp* gene is 610 bp while the 16S rDNA gene is 438 bp. PCR amplification process underwent two replicates to validate the results obtained (See Supplementary Table 1). A third screening was performed for selected individual samples that had conflicting results based on the two prior replicates. Therefore, the criteria set in reporting the certainty for *Wolbachia* infection is based on two successful amplification of the molecular markers. Amplified PCR products from each molecular marker were sent for sequencing to Eurofins, Operon – Tokyo.

### Identity of Wolbachia strains and their positions in phylogroups

All sequences were subjected to the Nucleotide Basic Local Alignment Search Tool (BLAST) and compared to deposited *Wolbachia* sequences in GENBANK. Next, selected sequences of *Wolbachia* strains (Table 1) and those obtained in the study underwent multiple alignment using Clustal W in MEGA 6^43^. After editing, the final length used for phylogenetic inference analyses was 398 bp and 732 bp for *wsp* and 16S rDNA respectively. The identities and relationships of the *Wolbachia* strains obtained in our study were determined by performing the Bayesian method in PhyML 3.0 software with 1000 bootstrap replicates^44^. The Smart Model Selection^45^ was also utilized to set the parameters for *wsp* as GTR+G (number of estimated parameters k = 232, Akaike Information Criterion (AIC) = 4897.31702) and 16S rDNA as GTR+G+1 (number of estimated parameters k = 207, Akaike Information Criterion (AIC) = 5332.88688). All sample sequences were submitted to GENBANK with Accession numbers____ - _____.

**Table 1.** Representative *Wolbachia* type sequences from different insect hosts in *wsp* and 16S rDNA molecular markers.

| **Molecular**<br>**Marker** | **Host** | ***Wolbachia***<br>**supergroup** | **Accession**<br>**Number** |
| --- | --- | --- | --- |
| wsp | *Drosophila melanogaster* | A | AF020072 |
|  | *Aedes albopictus* | A | AF020058 |
|  | *Glossina morsitans* | A | AF020079 |
|  | *Drosophila simulans* (Riverside) | A | AF020070 |
|  | *Muscidifurax uniraptor* | A | AF020071 |
|  | *Phlebotomus papatasi* | A | AF020082 |
|  | *Glossina austeni* | A | AF020077 |
|  | *Culex pipiens* | B | AF020061 |
|  | *Culex quinquefasciatus* | B | AF020060 |
|  | *Aedes albopictus* | B | AF020059 |
|  | *Ephesia cautella* | B | AF020076 |
|  | *Dirofilaria immitis* | C (Outgroup) | AJ252062 |
| 16S | *Nasonia longicornis* | A | M84691 |
|  | *Muscidifurax uniraptor* | A | L02882 |
|  | *Aedes albopictus* | B | KX155506 |
|  | *Culex pipiens* | B | X61768 |
|  | *Nasonia vitripennis* | B | M84686 |
|  | *Onchocera volvulus* | C | AF069069 |
|  | *Dirofilaria immitis* | C | Z49261 |
|  | *Litomosa westi* | D | AJ548801 |
|  | *Folsomia candida* | E | AF179630 |
|  | *Mansonella ozzardi* | F | AJ279034 |
|  | *Dipetalonema gracile* | J | AJ548802 |
|  | *Rickettsia sp.* | Outgroup | U11021 |

### Statistical Analysis

A Clark-Evans test was performed to determine if the spatial distribution of *Wolbachia*-positive mosquito samples from each molecular marker have a pattern of complete spatial randomness. The test uses the aggregation index (*R*) where a value of > 1 suggests an ordered distribution and a value of < 1 suggests clustering. This analysis was performed using R program version 3.3.5^46^under package *spatstat* ^46^

## RESULTS

### Detection of Wolbachia through wsp and its phylogeny

From a total of 672 adult *Ae. aegypti* screened, 113 (16.8%) individual adult mosquito samples are infected with *Wolbachia* using the *wsp* marker (Table 2). Based on the study’s criterion (See methods), only 17 samples demonstrated one successful amplification, thus excluding them for further analysis. In addition, female/male ratio is 0.82 (Table 2). All sequenced amplicons resulted in a high degree of similarity (>98.0%) with deposited *wsp* sequences in GENBANK. The spatial distribution showed that 60 (43.0%) sampling sites (Figure 1b) contained *Wolbachia* positive mosquitoes with 1 – 8 individuals. Further analysis showed that the distribution of *wsp*-positive mosquito samples was significantly clustered (R = 0.003,p < 0.001). Figure 2 and Figure S1 show the phylogeny of *Wolbachia* sequences based on *wsp* sequences. Majority of the sequences were found in supergroup B (n=84) while the remaining were clustered in supergroup A (n=29). Based on descending order of sample sizes, sample sequences in supergroup B were identical (>99.0%) to *Wolbachia* type strains from selected hosts of *Ae. albopictus* (*wAlbB*) (n= 51), *Cu. quinquefasciatus, Cu. pipiens* (*wPip*), *Ae. aegypti wMel* strain (n= 23) and *Ephestia cautella* (*wCau*) (n= 10). The sample sequences from supergroup A were either similar (98.0-99.0%) (n = 8) or identical (>99.0%) (n= 21) to the type strain (*wAlbA*) from host *Ae. albopictus*.

**Table 2.**
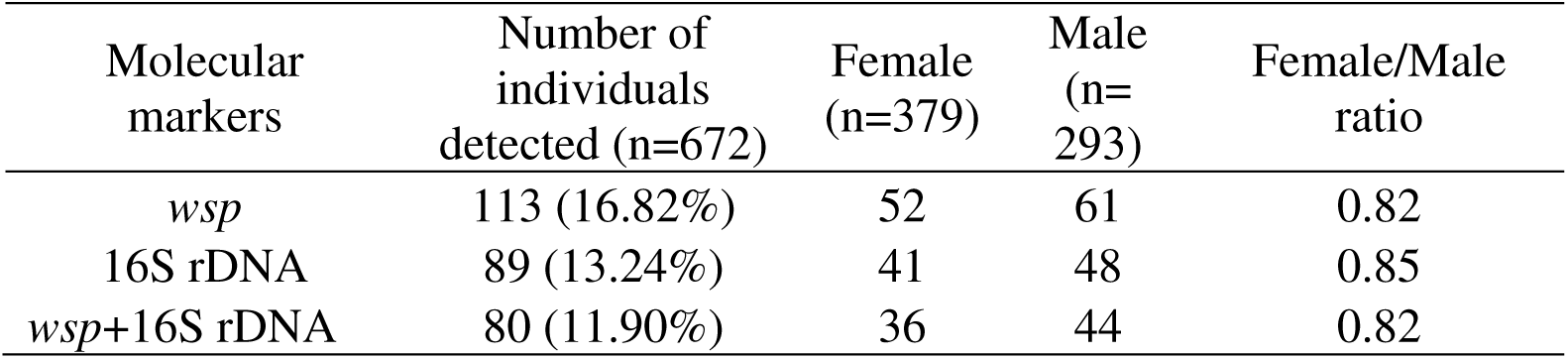
Summary *of wsp and 16S rDNA* detection results in *Ae. aegypti*

| Molecular markers | Number of individuals detected (n=672) | Female (n=379) | Male (n=293) | Female/Male ratio |
| --- | --- | --- | --- | --- |
| *wsp* | 113 (16.82%) | 52 | 61 | 0.82 |
| 16S rDNA | 89 (13.24%) | 41 | 48 | 0.85 |
| *wsp*+16S rDNA | 80 (11.90%) | 36 | 44 | 0.82 |

**Fig 2.**
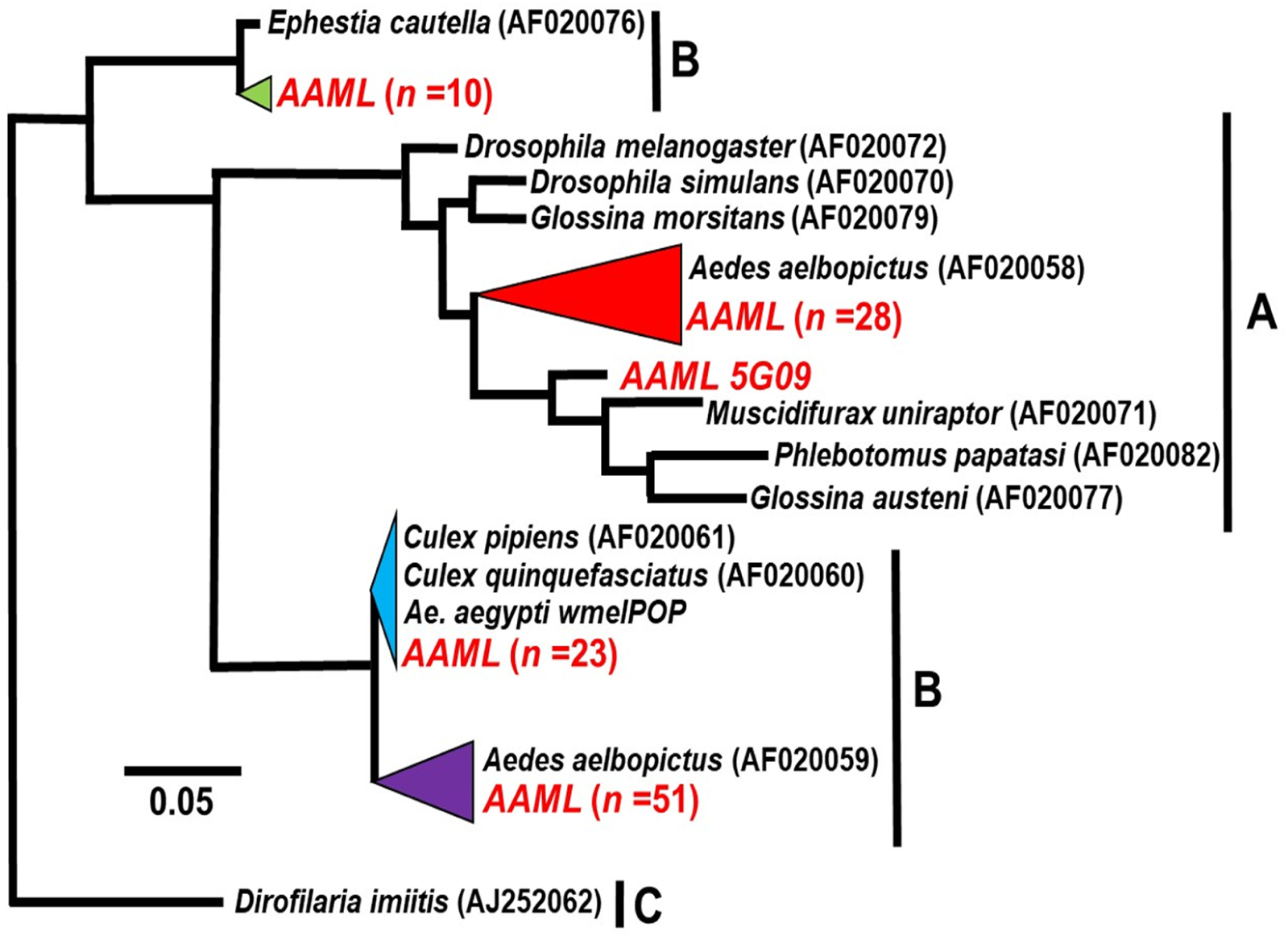
Phylogenic analysis of *wsp*. The alignment was analyzed in PhyML. Sample sequences of *Ae.aegypti* collected in Metropolitan Manila are in red, labeled as AAML (***<u>A</u>**e.* **<u>a</u>***egypti* **<u>M</u>**etropolitan Mani**<u>L</u>**a) and alphanumeric values indicate the unique code assigned to each *Ae. aegypti* individual sample. Merging (triangles) of sample and representative *Wolbachia* sequences was done to show degree of similarity (98-100%). Supergroups were indicated as A – C depending on the representative sequences used. The phylogenetic trees are re-drawn for better visualization, thus an expanded version can be viewed in Supplemental Figure S1. Please refer to Table 1 for the *Wolbachia* type sequences (ingroup and outgroup) for both markers.

### Detection of Wolbachia through 16S rDNA and its phylogeny

For 16S rDNA, 89 (13.2%) individual adult mosquito samples were infected with *Wolbachia* (Table 2). 20 individual mosquito samples generated one successful 16S rDNA amplification, thus, excluding them for further analysis. Furthermore, female/male ratio is 0.85 (Table 2). 50 (36.0%) sampling sites (Figure 1c) contained *Wolbachia*-positive mosquitoes ranging from 1-8 individuals and the distribution of 16SrDNA-positive individuals revealed to be clustered or aggregated (R = 0.001,p < 0.001). All sequenced amplicons resulted in a high degree of similarity (>98%) with deposited 16S rDNA Wolbachia sequences in GENBANK. Nearly all 16S rDNA sample sequences (n=85) (Figure 3, Figure S2) were grouped in supergroup B. Only one sample sequence was identical to *Nasonia vitripennis* while the remaining sample sequences were up to 99% similar from the selected hosts of the supergroup. The remaining sample sequences (n=4) were grouped in supergroup C & J. One sample sequence was highly similar (>99%) with *Dirofilaria immitis* while the remaining were 98-99% similar from the selected hosts of the supergroup.

**Fig 3.**
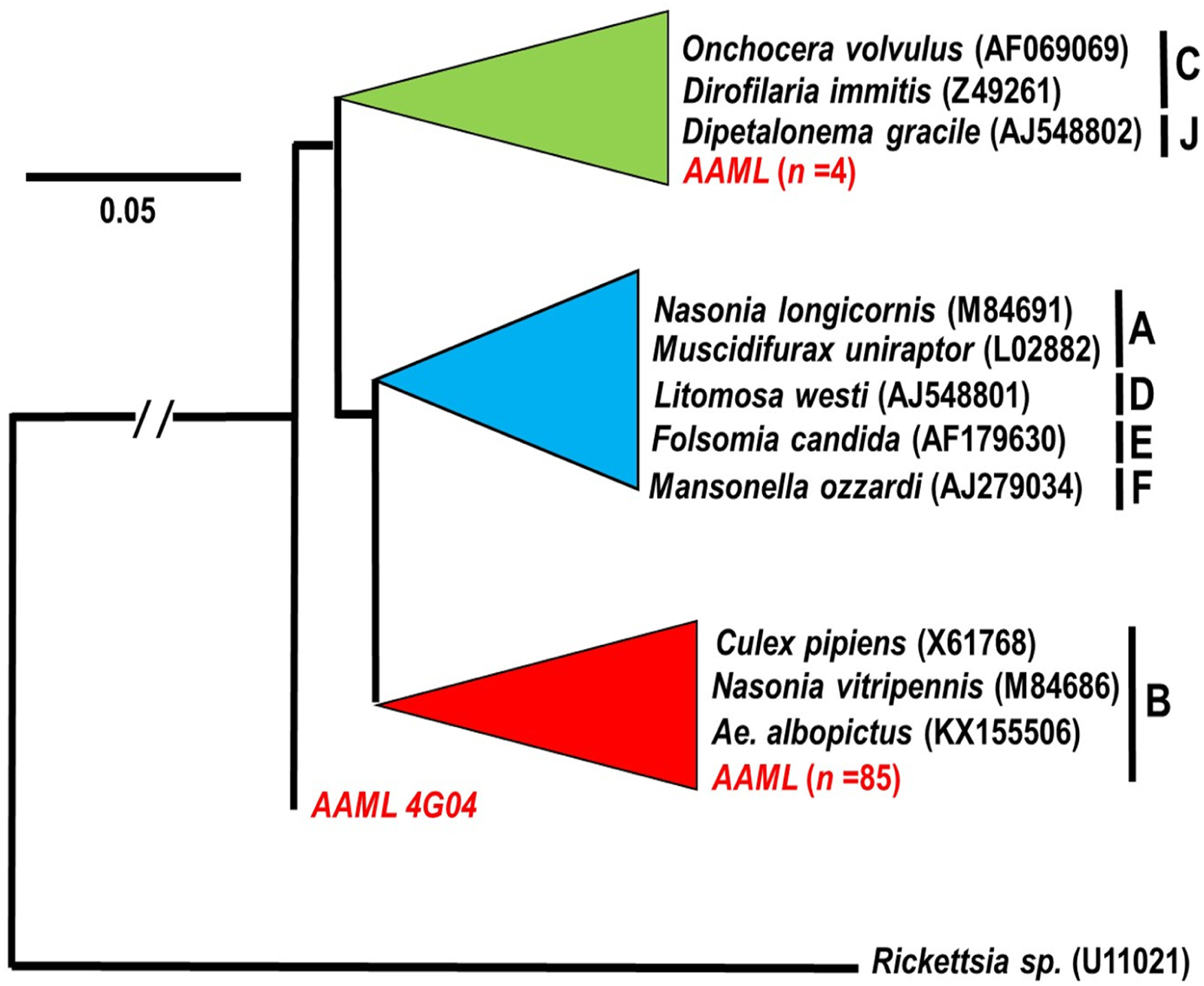
Phylogenic analysis of 16S rDNA. The alignment was analyzed in PhyML. Sample sequences of *Ae.aegypti* collected in Metropolitan Manila are in red, labeled as AAML (***<u>A</u>**e.* **<u>a</u>***egypti* **<u>M</u>**etropolitan Mani**<u>L</u>**a) and alphanumeric values indicate the unique code assigned to each *Ae. aegypti* individual sample. Merging (triangles) of sample and representative *Wolbachia* sequences was done to show degree of similarity (98-100%). Supergroups were indicated as A – J depending on the representative sequences used. The phylogenetic trees are re-drawn for better visualization, thus an expanded version can be viewed in Supplemental Figure S2. Please refer to Table 1 for the *Wolbachia* type sequences (ingroup and outgroup) for both markers.

### Comparison of 16S rDNA and wsp for Wolbachia detection and phylogeny

From the 113 and 89 positively detected mosquito individuals from *wsp* and 16S rDNA respectively, 80 (11.90%) individual samples yielded positive amplification in both markers (Table 2). In *wsp* positive detection (n=113), 80 had two successful amplification of the 16S rDNA marker while 27 had only one amplification of 16S rDNA and the remaining 6 had no successful amplification on 16S rDNA marker. On the other hand, the 89 individual samples deemed 16S rDNA positive for *Wolbachia* showed 80 individuals had two successful amplification of the *wsp* marker while 9 had only one successful amplification on the said marker. Next, we focus on the supergroup classification of the 80 individual samples based on the *wsp* and 16S rDNA phylogeny. It was shown that 55 (69%) had the same classification in supergroup B while the remaining 25 (31%) showed a disparity in supergroup classification. Such difference, for example, showed that *wsp* identified the individual sample as supergroup A, but 16S rDNA reveals to be either supergroup B or C & J.

## DISCUSSION

Our study was able to demonstrate the detection of the endosymbiont *Wolbachia* in field-caught adult *Ae. aegypti*. Notably, the main reason for the positive detection, especially in *wsp*, is because of the procedural modifications or optimization in the amplification of the said marker. A case in point, for example, why optimization is necessary is the evidence presented in the malaria mosquito vector, *An*. *gambiae*. Previous studies had reported no observed natural *Wolbachia* infection in this mosquito vector^26-31^; however, the endosymbiont was successfully detected in *An. gambiae* from Burkina Faso, West Africa using an optimized *wsp* protocol^32,33^. Another potential reason for a positive detection was the study’s sample size. Based on several literature on assessing the prevalence of *Wolbachia* in different mosquito species, the highest number of *Ae. aegypti* individuals screened was 119^30^which resulted in non-detection of the endosymbiont. As compared to the actual study (n= 672), the sample sizes from previous studies were low; thus, larger sample size would provide a more accurate estimate of the prevalence of *Wolbachia* infection. Similarly, these reasons were clearly emphasized by recent studies on why earlier investigations may have underestimated the actual incidence of *Wolbachia* infection from different insect hosts^48,49^.

Our study acknowledges the uncertainties associated with conventional PCR detection such as high false positive detection rates. With this in mind, the study was cautious in affirming a positive infection in each *Ae. aegypti* adult sample. First, the selection of markers is based on the recommendation of Simoes et al.^42^that two of its preferred primer sets (e.g. *Wspecf* and *Wspecr*) was determined to produce the lowest false positive and false negative rates. Secondly, our study performed replications with a stringent criterion for a successful *Wolbachia* infection on each mosquito sample. Although there are several genetic markers (e.g. MLST genes) and techniques (e.g. IFA, FISH or whole-genome sequencing) available, the primary intention of this study is to detect *Wolbachia* infection in *Ae. aegypti* initially using this PCR-Based approach.

Linking our findings with the previous studies^35-37^which reported *Wolbachia* in *Ae. aegypti* may incidentally provide a clear picture of its infection status. First, the probable density of the endosymbiont found in this mosquito vector may be low. Even though our study did not measure the actual density, a 40-cycle PCR amplification procedure or a long PCR run^50^may detect a small amount of *Wolbachia* present. It partly supports the results presented from metabarcoding studies^36,37^where a low number (2-4) of *Wolbachia* sequence reads were detected in both the larvae and adult *Ae. aegypti* mosquito. These can be another potential reason why earlier prevalence studies were not able to detect *Wolbachia* in *Ae. aegypti* samples. Moreover, the low probable density of the endosymbiont may also translate to the observed low infection rate (13-16%) found in our study. This again partly supports metabarcoding studies^36,37^where only two *Ae. aegypti* mosquito pools had the presence of these low number *Wolbachia* sequences. On the other hand, our results are in contrast with the report from *Ae. aegypti* larvae (n=16 individuals) in Malaysia which resulted in a 50% infection rate^35^. However, there could be some uncertainties to this estimate because of its small sample size and, more importantly, the collected larval samples may be siblings from the same female *Ae. aegypti* mosquito. The limitation as mentioned earlier prompted us to conduct an individual-based adult mosquito detection so that it can present a better and explicit estimation of the infection rate. Secondly, we assume that the *Wolbachia* strain/s found in *Ae. aegypti* can be maternally-inherited due to the following reasons: (a) reported positive infections in larval samples from the previous studies^35-37^and (b) detecting positive infections in male *Ae. aegypti* mosquitoes (our study, Table 2). However, there is still a need to present direct evidence of maternal transmission of this endosymbiont during the developmental stages of *Ae. aegypti* since all studies, including ours, were performed independently.

Lastly, the *Wolbachia* strains infecting *Ae. aegypti* have been shown in our study belong to supergroups A and B. Both *wsp* and 16S rDNA phylogeny showed that majority of the individual samples belong to supergroup B while a small number of individual samples were found in supergroup A (based on *wsp*). Detecting different *Wolbachia* strains in a single mosquito species is relatively common especially in medically important mosquitoes, *Ae. albopictus*^51,52^ and *An. gambiae*^32^, and other insect host species (e.g. *Drosophila* species^51^). Since our study presented a majority of our sample sequences belonging to supergroup B, this was also the same observation reported by previous studies^35-37^. Dipterans, especially mosquitoes, are commonly infected by these *Wolbachia* strains from supergroups, A and B. It has been shown to cause parasitism towards its insect host by producing phenotype effects such as cytoplasmic incompatibility, male killing, and feminization^11,53^. Nevertheless, whether the identified *Wolbachia* strains in *Ae. aegypti* possess these phenotypic effects remains unclear. Also, further studies are needed to ascertain the pathogenic impact of this local endosymbiont to the mosquito vector. More importantly, it is very essential to determine whether these identified *Wolbachia* strains could render *Ae. aegypti* a less effective vector by blocking key arboviruses such as dengue. It is also worth mentioning that some individual samples have shown to be similar with *Wolbachia* strains found in supergroups C and J based on 16S rDNA. These two supergroups are not generally found in dipterans especially in mosquitoes. It is likely that our 16S rDNA amplified the *Wolbachia* strain residing in the roundworm, *Dirofilaria immitis*. *Ae. aegpyti* mosquitoes are also known to carry this parasitic nematode to certain mammals, such as dogs^54^. This observation was also reported in one of the metabarcoding studies^37^that showed sequences of *Wolbachia* from *Dirofilaria immitis*. However, when these 16S rDNA results were compared to the *wsp* results in our study, it showed the *Wolbachia wsp* sample sequence of the same mosquito individuals belong to supergroup B. We can only infer that the inconsistent results observed in our study may stem towards the sensitivity and specificity of the markers used. The *wsp* gene marker has been likened to antigen protein typing in screening pathogenic bacteria where it can be a perfect diagnostic tool for detecting *Wolbachia* infection^55,56^. However, it is unsuitable for phylogenetic analysis or deeper taxonomic relationship because of its extensive recombination and strong diversifying selection^11,57,58^. 16S rDNA, on the other hand, is known to be a conserved gene highly suited in bacterial identification and phylogeny, but its use in detecting *Wolbachia* infection has demonstrated varying results depending on the specific 16S rDNA primers^42^. It was emphasized that “no single protocol” can ultimately ensure the specificity and accuracy of 16S rDNA to detect *Wolbachia* infection^56^. Thus, further claiming that 16S rDNA markers in *Wolbachia* detection may be far from optimal^56^.

We consider our findings to be crucially important especially if the Philippines would implement or approve two scenarios in the release of: (a) *Wolbachia*-infected (e.g. *w*MelPop or *w*Mel) mosquitoes or (b) local *Wolbachia* strains found by our study in dengue-endemic areas. In the first scenario, a vital consideration is the presence of “bidirectional incompatibility” mechanism between the intended *Wolbachia* strain (e.g. *w*MelPop or *w*Mel) to be released and the present local strain found in the mosquito. There are instances that two strains in one host cannot stably coexist with each other because the naturally occurring strain is preventing the intended strain to reach fixation or establishment^59-61^. It would serve as an impediment to the intentional spread of *Wolbachia* strain to the mosquito population. It was suggested that to overcome this incompatibility is to remove the existing natural strain inhabiting the mosquito vector or to perform a “superinfection” where the intended *Wolbachia* strain induces unidirectional incompatibility with the natural strain^62^. Nevertheless, it very important to re-examine the infection status of *Wolbachia* in *Ae. aegypti* mosquitoes in intended areas prior a mass release program. If the second scenario, utilizing the release of local *Wolbachia* strains, is implemented, there are specific considerations that should be addressed for a successful population replacement. The first and most important consideration is to determine whether these local strains may exhibit the same phenotypic effects and pathogen blocking of *w*Mel strain to *Ae. aegypti*. Currently, these characteristics are still unknown and therefore crucial if utilized for mass release. Another consideration is endosymbiont’s density in the mosquito vector. Mosquito species naturally infected with *Wolbachia* are not ideal candidates due to the changing molecular interactions between *Wolbachia* and the host over time^63^. The result of this symbiosis is the amount of bacterial density found in the mosquito host where it can influence the intensity of *Wolbachia*-induced phenotypic or anti-viral effects^22,62,64,65^. Newer infections (e.g. tansinfections) are shown to produce high bacterial density while natural infections lead to lower bacterial density due to the adaptation of the host to the endosymbiont infection over time. In our study, we infer that the local *Wolbachia* strains are in low density inside its host, *Ae. aegypti*. If this is the case, it will result in a reduced physiological and anti-viral impact of the strain to the mosquito vector. However, high *Wolbachia* density which also possesses strong inhibitory effects against insect viruses had been observed from natural *Wolbachia* strains with a long-term association from its host^66,67^. The last consideration is the low infection rate. It raises the question, more importantly to the population replacement approach, if any of the local *Wolbachia* strains could be sustained for an extended period or possess the ability to infect the mosquito population thoroughly. Studies had suggested that a successful strain used in population replacement or invasion should reach an infection rate of >90% and should remain at this rate over an extended period of time^68-70^. Thus, utmost consideration in the infection status of *Wolbachia* and its role in *Ae. aegypti* is necessary for a *Wolbachia-*based vector control program to be successful, efficient and, as well as, effective.

## ACKNOWLEDGEMENTS

We would like to thank M.J.L.B. Martinez, J.D.R. Capistrano, V.S.P. Tiopianco, B.M.C Orantia, C.R. Estrada, M.G. Cuenca, K.M. Viacrusis and L.F.T. Hernandez for their valuable work in the collection of the mosquitoes. Also we are grateful to the valuable and pertinent comments of the anonymous reviewers. This work is funded by the JSPS Grant-in-Aid for Scientific Research (16H05750, 17H01624, 17K18906), JSPS Bilateral Joint Research Projects, and Leading Academia in Marine and Environmental Pollution Research – Ehime University (Y29-1-8)

## AUTHOR CONTRIBUTIONS

T.M.C., D.M.A. and K.W. designed the experiments. T.M.C., K.H., and R.K.H. performed the experiments. T.M.C., K.H., and R.K.H. performed the sequencing while T.M.C., K.W. and D.M.A. accomplished the phylogenetic analysis. T.M.C. wrote the manuscript along with and K.W. All authors reviewed the manuscript and approved on its submission.

## DATA AVAILABILITY

Demographic profiles (location and sex) and detection status from each individual *Ae. aegypti* adult mosquito used in the study are presented in the Supplementary. Accession numbers of Nucleotide sequences of PCR-amplified fragments of *wsp* and *16S* have been deposited in the GENBANK nucleotide database under accession numbers ______to ______and _____to ______respectively.

## SUPPLMENTAL MATERIAL

**Table S1.** Demographic profile (Sex, Sampling Site Code, Location), Detection status (*wsp* and 16S rDNA) of all individual adult *Aedes aegypti* mosquitoes used in the study. Positive *Wolbachia* infection in mosquito samples presents the supergroup classification and GENBANK accession number.

**Figure S1.** Complete *wsp* phylogeny of *Wolbachia* from *Ae. aegypti* (n=113). The alignment was analyzed in the program PHYML and *Wolbachia* host *Dirofilaria immitis* was selected as an outgroup. All sample sequences are indicated in red dots. The condensed version of this tree is presented as Figure 1.

**Figure S2.** Complete *16S* rDNA phylogeny of *Wolbachia* from *Ae. aegypti* (n=85). The alignment was analyzed in the program PHYML and *Rickettsia sp.* was selected as an outgroup. All sample sequences are indicated in red dots. The condensed version of this tree is presented as Figure 2

